# Tandem Walk in Simulated Martian Gravity and Visual Environment

**DOI:** 10.1101/2022.02.01.478711

**Authors:** Marissa J. Rosenberg, Matthew Koslovsky, Matthew Noyes, Millard F. Reschke, Gilles Clément

**Affiliations:** KBR, Houston, TX, USA; Department of Statistics, Rice University, Houston, TX, USA; Neuroscience Laboratory, NASA Johnson Space Center, Houston, TX, USA; UNICAEN-INSERM U1075 COMETE, University of Caen Normandy, Caen, France

**Keywords:** Spatial disorientation, Virtual reality, Martian gravity, Tandem walk

## Abstract

Astronauts returning from long-duration spaceflights experience visual-vestibular conflicts that causes motion sickness, perceptions that the environment is moving when it is not, problems with walking, and other functional tasks. To evaluate whether astronauts will have similar decrements associated with visual-vestibular conflicts after they land on Mars following exposure to weightlessness, participants were held by a device that offloads their weight; first entirely (0 G) for 10 minutes, and then partially (0.38 G) or not at all (1 G) for 15 minutes. Tandem (heel-to-toe) walk was used to assess the subjects walking performance. Ten subjects performed 2 trials of 10 steps on a medium-density foam surface. Four conditions were investigated: (a) 1 G in virtual reality (VR); (b) 1 G in VR with a superimposed disorienting optokinetic simulation (VR+DOS); (c) 0.38 G in VR; and (d) 0.38 G in VR+DOS. Tandem walk performance decreased in VR+DOS compared to VR in both 1 G and simulated 0.38 G. Tandem walking performance in VR+DOS was better in 0.38 G compared to 1 G. Tandem walking performance in VR+DOS in 1 G was not significantly different from tandem walking performance after spaceflight or bed rest. The increased tandem walking performance in 0.38 G compared to 1 G was presumably due to an increased cone of stability, allowing larger amplitude of body sway without resulting in a fall. Tandem walking on a compliant foam surface in VR+DOS is a potential analog for simulating postflight dynamic balance deficits in astronauts.

## INTRODUCTION

Visual-vestibular alterations occur during critical periods of space travel, such as during entry into weightlessness and during return to Earth’s gravity. Disturbances induced through visual-vestibular conflict typically include motion sickness, perceptual illusions, changes in eye-head coordination, and alteration in balance and gait after landing. The longer astronauts are exposed to microgravity, the more intensely they experience these disturbances. This pattern poses a significant challenge for astronauts who participate in future long-duration exploration missions to the Moon or Mars. The current scenario for a human Mars mission includes an 8-month journey in weightlessness (0 G) before landing. It is therefore critical to assess how humans will perform in martian gravity after they have adapted to 0G (Paloski et al., 2008).

During reentry into a gravitational field, astronauts commonly report exaggerated motion of their surroundings when they move their head (Reschke et al., 2017). While performing head movements in pitch, several crewmembers reported a perceived translation of the visual surroundings and the sensation that they were accelerating faster than they actually were. Also, their perceived self-translation and perceived head roll were greater in rate and displacement amplitude than the actual head roll. The displacement amplitude of yaw head movements made immediately after landing also felt exaggerated (Clément & Reschke 2008).

Returning crewmembers also experience difficulties standing and walking, especially with their eyes closed. They experience postural instability and walk slowly even after missions of relatively short duration, whereas spaceflight-induced changes in their in muscular strength are minimal due to the efficacy in-flight exercise countermeasures (Bloomberg et al., 1997; Black & Paloski, 1998). Thus, postural changes cannot be solely ascribed to spaceflight-induced skeletal muscle changes. It appears that some of these deficits are due to gaze destabilization (oscillopsia) because of a reduced ability to engage in compensatory head pitch movements during locomotion (Peters et al., 2011). The organization of astronauts’ gait pattern is only slightly disrupted after spaceflight, which indicates that they rely significantly on visual feedback when walking after return from space since their vestibular system is decidedly impaired.

During the Apollo missions, astronauts walked slower during extra-vehicular activities on the lunar surface than they did on Earth (Berry & Homick 1973). However, falls and near falls were frequent due to the ruggedness of the terrain, limited mobility and visibility in the suit, and astronauts automatic postural reactions did not correct the disequilibrium. Indeed, the automatic postural reactions in response to tipping or stumbling occurred too fast in lunar gravity (0.16 G), so these reactions caused further disruption to astronauts’ balance (Kubis et al., 1972).

Studying the visual-vestibular conflict that occurs through adaptation of the vestibular system in weightlessness is difficult to induce in a normal 1 G environment. Thus, researchers have not been able to directly study the capability of humans to walk in martian gravity (0.38 G). Rather than modifying the gain of the vestibular system, experimental protocols frequently rely on modifying visual input to induce visual-vestibular conflict. Ground-based simulations using suspension devices have been used to predict the mechanical work and metabolic expenditures required for walking and running in martian gravity (Cavagna at al., 1998), but no one has studied the relative contribution of visual and somatosensory cues to dynamic balance in martian gravity.

The first objective of this study was to investigate how exaggerated motion of visual surroundings affects dynamic balance in simulated martian gravity. The second objective was to simulate the walking deficits seen in astronauts after spaceflight for evaluating potential countermeasures. A virtual reality environment simulated the astronauts’ postflight illusory movements caused by head movements. A suspension device was issued to simulate weightlessness (0 G) and martian gravity (0.38 G). The Tandem Walking Test was used to assess dynamic balance control in this simulated visual and gravitational environment. We chose the Tandem Walking Test because it is used in clinics to assess patients who may be at risk for falls (Cohen et al., 2012); it has also been used to evaluate dynamic balance control in astronauts returning from spaceflight (Homick & Reschke 1977; Reschke et al., 2017; Miller et al., 2018; Mulavara et al., 2018). The third objective of this study was to compare the tandem walk performance in simulated weightlessness and martian gravity in our study with previously collected data during tandem walk on a hard floor with the eyes closed astronauts after spaceflight and in subjects and after bed rest (Miller et al., 2018; Mulavara et al., 2018).

## METHODS

### Subjects

Ten healthy volunteers (5 male, 5 female; aged 32.3 ± 5.3 years) participated in this study. Subject were selected through NASA’s Test Subject Screening procedures. All subjects passed a Class III flight physical. This study was carried out in accordance with the recommendations of the NASA Johnson Space Center Institutional Review Board and were performed in accordance with the ethical standards established by the 1964 Declaration of Helsinki. All subjects provided written, informed consent before participating in the study. The protocol was approved by the NASA Johnson Space Center Institutional Review Board. All tests were conducted at the NASA Johnson Space Center, Houston, TX, USA.

### Active Response Gravity Offload System (ARGOS)

The NASA ARGOS system allows human subjects to perform functional tasks in simulated reduced gravity (Fig. 1). Subjects can move freely inside a 12.5-m long x 7.3-m wide x 7.3-m high steel structure while in a harness suspended by a steel cable. ARGOS uses robotic mechanisms and computer controlled electric motors to supply a continuous vertical offload of 0% (1 G) to 100% (0 G) of a person’s weight during walking, running, and jumping. Rotational motion is also accommodated by various gimbal interface mechanisms (Cunningham, 2010). For this study, ARGOS was utilized to off-load 62% of a person’s weight to enable testing in a 0.38 G setting.

**Figure 1.**
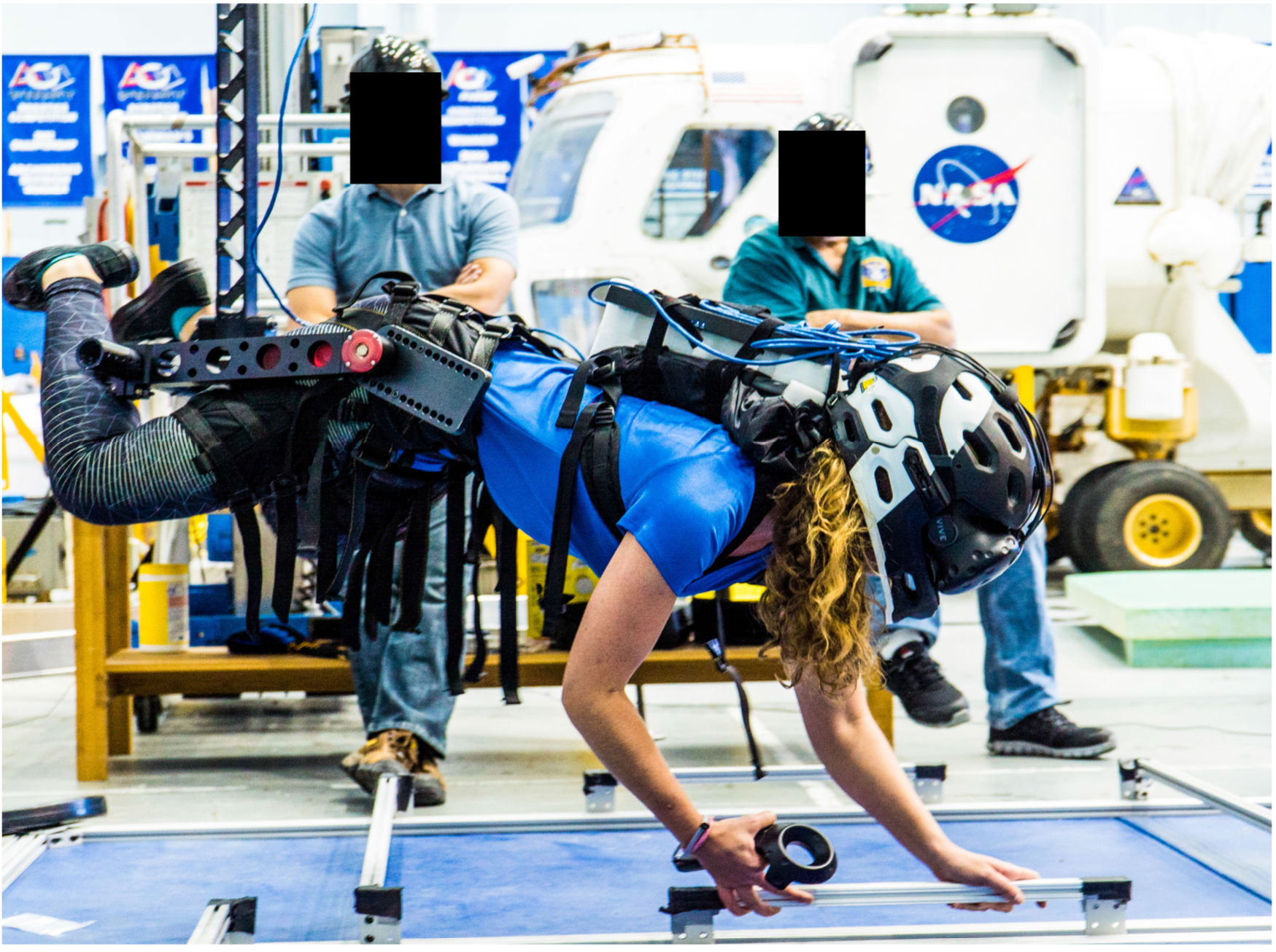
NASA’s Active Response Gravity Offload System (ARGOS) is used to offload 100% of the subject’s body weight (simulated 0 G). The subject is viewing handholds in virtual reality and using physical handholds to move herself around the environment.

### Virtual Reality (VR)

A virtual reality headset (Vive, HTC Corporation, New Taipei City, Taiwan) was used to visualize the International Space Station (ISS) mockup. Subjects could: (a) explore the exterior of the ISS (we refer to this as EVA for extra-vehicular activity) using hand rails to ambulate; (b) watch the Earth rotate below; and (c) look at the stars in the sky. Although the subjects’ feet, body weight, and vestibular system were all experiencing 1 G when interacting with the ISS (i.e. ambulating along the hand rails), the law of physics for the visual motion were the same as they would be in actual weightlessness. This VR simulation of the ISS while in 1 G condition primed subjects in VR before testing functional task performance with VR+DOS.

The subjects also wore the virtual reality headset while their body was 100% unloaded (simulated 0 G) using ARGOS (Fig. 1). Subjects performed various intra-vehicular activities (IVA), such as moving inside the various ISS modules and drilling using a virtual tool controlled by a hand controller. These tasks were designed to encourage use of their hands rather than their feet to move themselves and to perform tasks in a “floating” position, as is the case in a true 0 G environment. VR simulation of the ISS while in 100% unloaded condition primed subjects before making the transition to partial gravity, as would occur in an actual Mars mission landing.

### Disorienting Optokinetic Stimulation (DOS)

A disorienting optokinetic stimulation (DOS) was superimposed onto a scene of the martian environment (VR+DOS) (Fig. 2). The DOS was composed of a checkerboard pattern that moved in roll and yaw in the same direction and velocity as the subject’s head movements. The objective was to mimic the perceptual illusions reported by astronauts when they move their head after spaceflight (Clément & Reschke 2018). When turning the head to the right or left, subjects had the feeling of exaggerated translation to the right or left, respectively.

**Figure 2.**
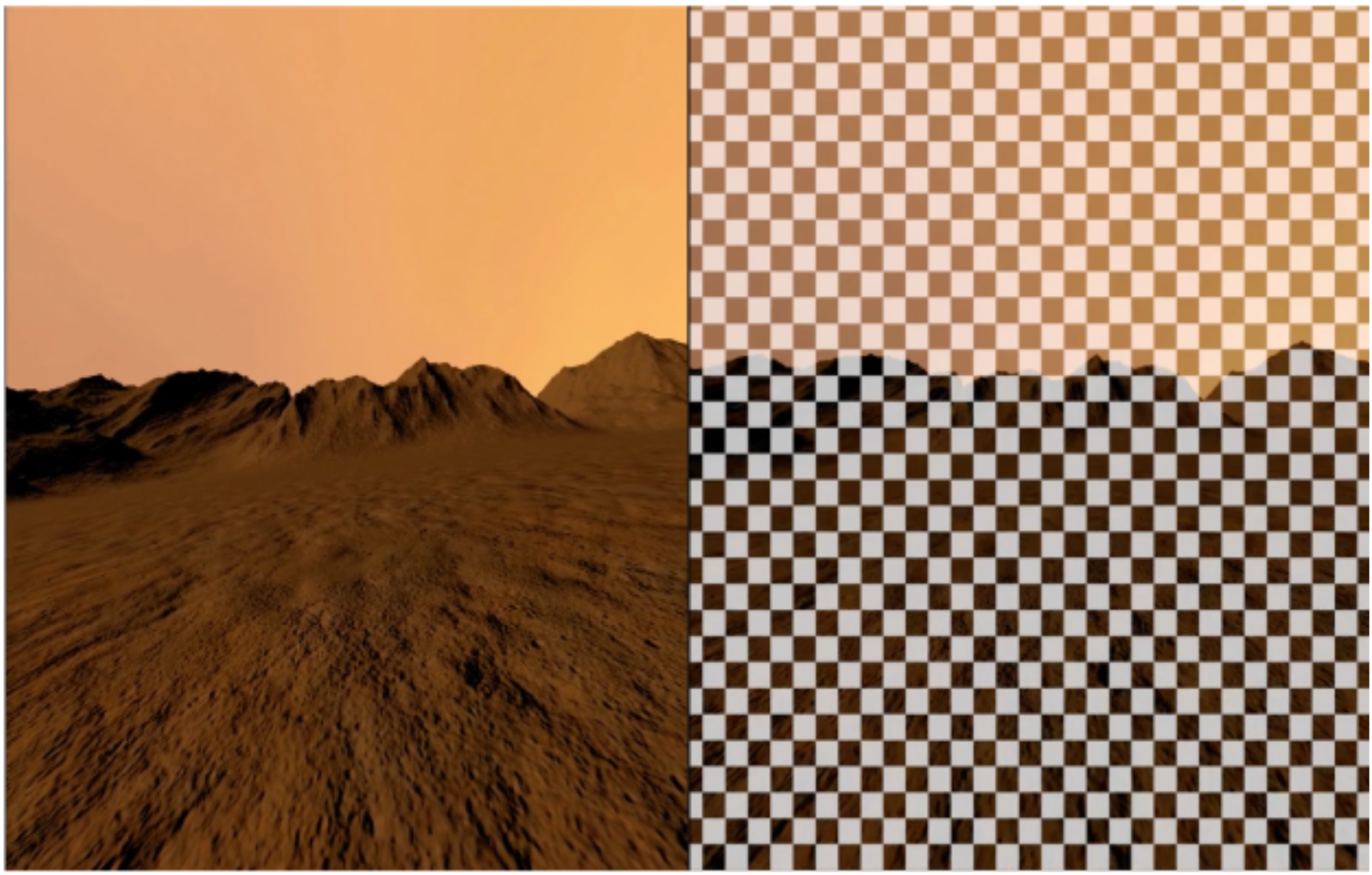
Left: Virtual scene of a martian environment (VR). Right: The virtual martian scene with the optokinetic stimulation shading layer superimposed (VR+DOS).

### Tandem Walk

Subjects were asked to walk heal-to-toe for 10 steps at a self-selected pace with their arms crossed over their chest. Subjects were then asked to turn 180 degrees and perform another 10-step tandem walk. The laboratory floor was covered by a 10-cm-thick medium density foam (Sunmate Foam, Dynamic Systems, Inc., Leicester, NC, USA) to make proprioceptive information (i.e. ankle proprioception) less reliable when walking, as is the case after adaptation to 0 G (Miller et al., 2018; Mulavara et al., 2018). During the Tandem Walking Test, subjects wore the virtual reality headset and were attached to the ARGOS suspension system (see below) (Fig. 3).

**Figure 3.**
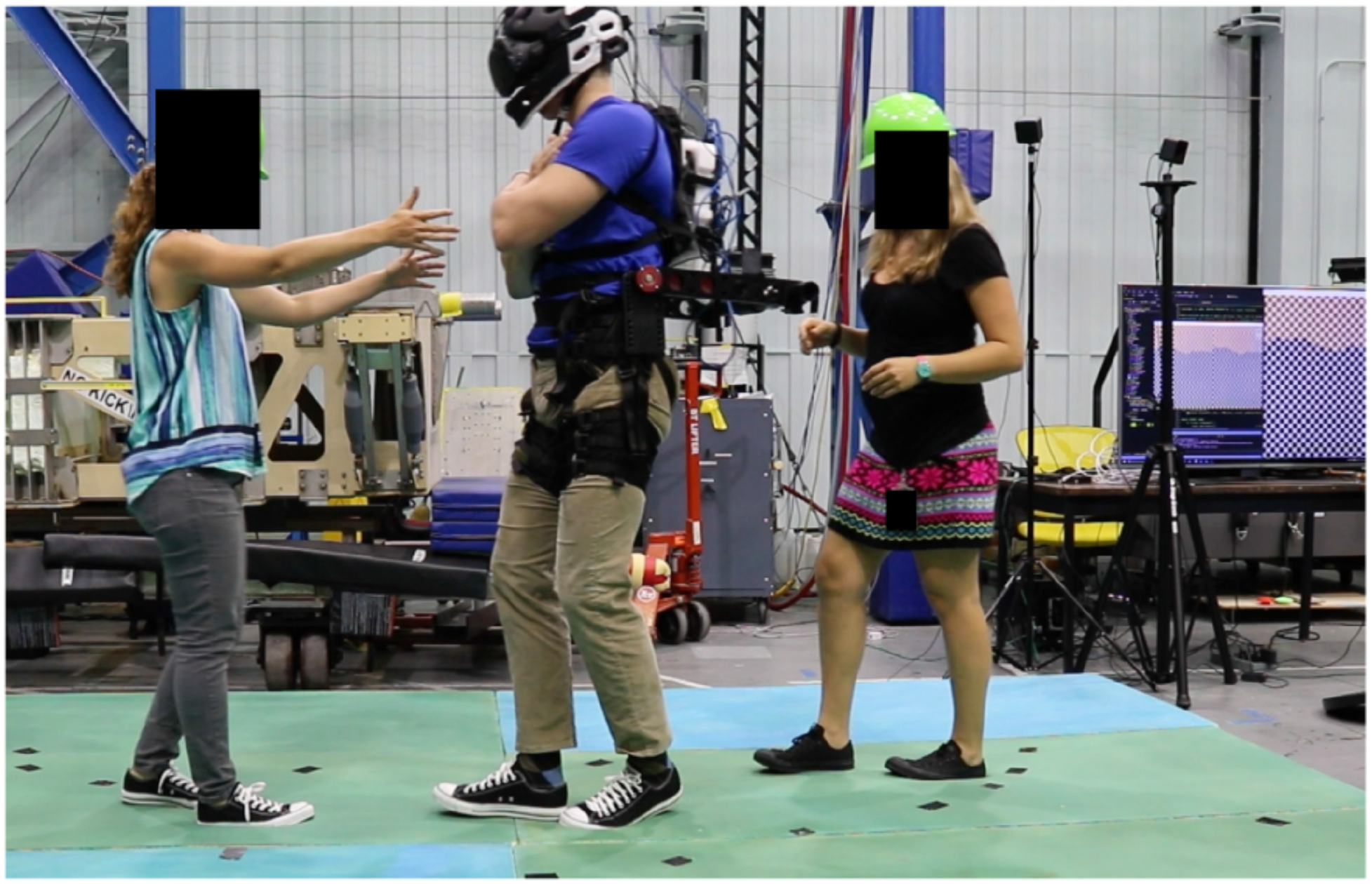
A subject, assisted by 2 operators, is performing the Tandem Walking Test on medium density foam with 62% of their body weight offload (simulated 0.38 G) while visualizing a martian environment. The subject and operators gave written informed consent for the publication of this image.

Each subject was tested during 4 separate sessions that included two spaceflight activities (Extra-vehicular activity, EVA; intra-vehicular activity, IVA), two visual conditions (VR, VR+DOS), and three body weights (equivalent to 1 G, 0 G, and 0.38 G) (Fig. 4). The protocol for each session was as follows: (a) 10 minutes of EVA or IVA activities in 1 G and simulated 0 G, respectively, while viewing a virtual mockup of the ISS; (b) a 2-minute transition to the foam surface with eyes closed, followed by approximately 3 minutes to adjust the virtual scene of Mars and body weight; (c) a 5-minute familiarization session while subjects performed a series of tasks while viewing a martian environment; and (d) the two 20-step tandem walks with VR or with VR+DOS. The test sessions were separated by an average of 4.3 days and the order of sessions was randomized.

**Figure 4.**
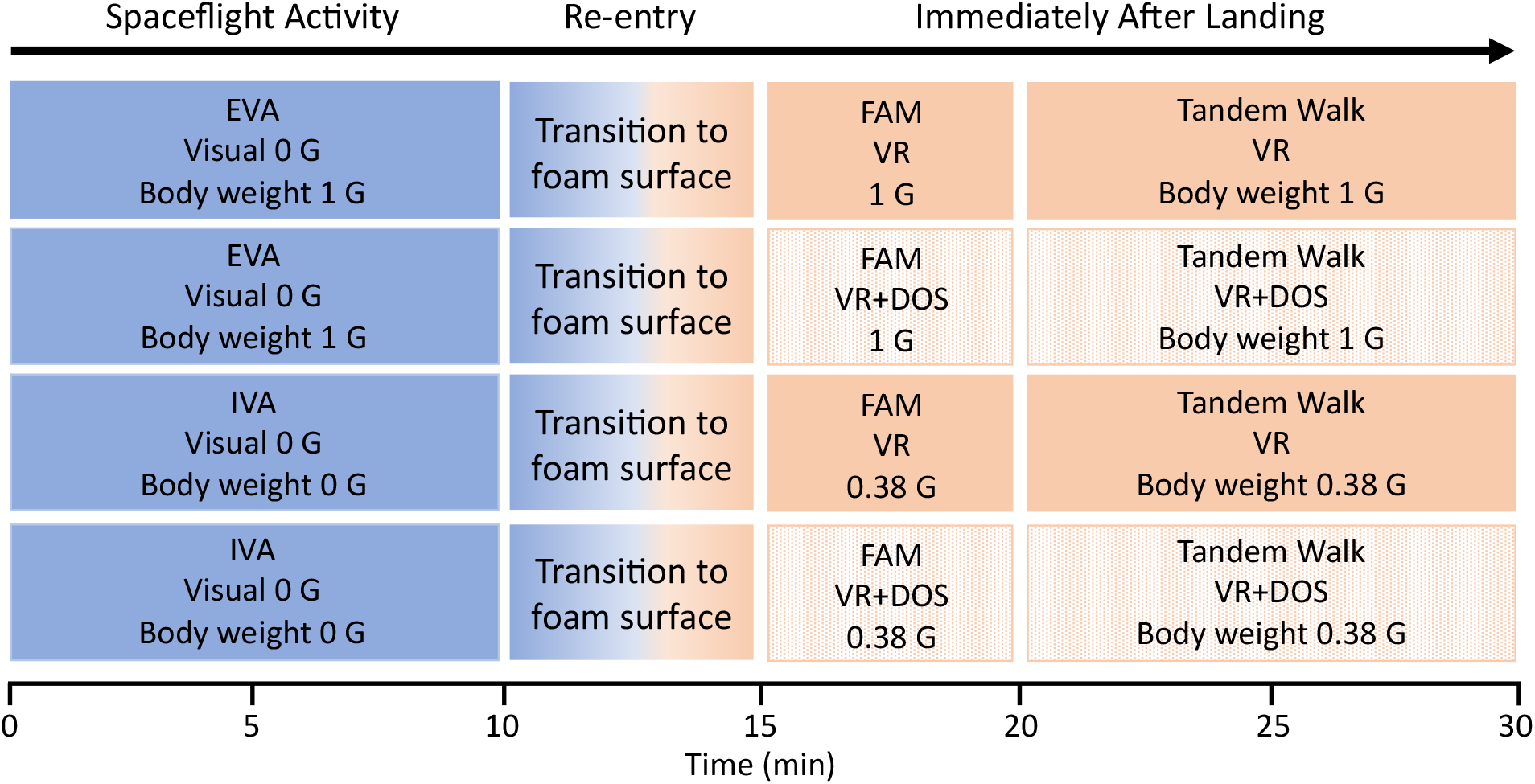
Diagram showing the conditions during the 4 test sessions. First the subjects performed extra-or intra-vehicular activities (EVA, IVA) while visualizing the ISS in virtual reality (Visual 0 G) with the body unloaded 0% (1 G) or 100% (0 G). Second, the subjects transitioned to a foam surface, and body unloading simulated the terrestrial (1 G) or martian (0.38 G) gravitational field. Third, the subjects were familiarized (FAM) with the visual environment and the body unloading conditions. Fourth, the subjects performed the Tandem Walking Test in 4 conditions: 1 G in VR; 1 G in VR+DOS; 0.38 G in VR; 0.38 G in VR+DOS. The order of sessions was randomized across subjects.

### Data Analysis

The video of each trial was recorded. Three reviewers examined the videos independently to determine the number of correct steps during each trial. A ‘‘misstep’’ was defined as any of the following: (a) the subject’s stepping foot crossing over the plant foot; (b) the subject stepping to the side before completing the step; (c) the subject’s stepping foot swinging in a wide, arcing path before stepping down; (d) a step duration greater than 3 s; or (e) a gap larger than 10 cm between the heel of the front foot and toe of the back foot when the step was completed (Miller et al., 2018; Mulavara et al., 2018). Videos for all trials across all sessions for a given subject were pooled, then the order was randomized to minimize reviewer bias based on their awareness of the session. After all reviewers completed their assessments, the median value was used to determine the Percent of Correct Steps for that trial.Teh Percent of Correct Steps for both trials were then averaged.

One subject in ARGOS was unable to bring their legs close enough together to walk heel-to-toe. The feet of another subject were not visible in the recorded video. Thus, we were unable to assess tandem walk performance for two subjects in 0.38 G and VR. For the remaining 8 subjects, repeated-measures analysis of variance (ANOVA) and post hoc t-tests were used to compare the Percent of Correct Steps between VR and VR+DOS in 1 G and 0.38 G.

In order to assess the validatity of using VR+DOS as an analogue for post flight tandem walk performance, we compared the Percent of Correct Steps during tandem walk on a foam surface in our study with the mean Percent of Correct Steps previously collected during tandem walk on a hard floor with the eyes closed in 13 astronauts after a 6-month stays on board the International Space Station (ISS) (Mulavara et al., 2018), in 7 astronauts after a 2-week spaceflight on board the Space Shuttle (STS) (Miller et al., 2018), and in 10 subjects after a 70-day bed rest in 6-deg head down tilt (Mulavara et al., 2018). For this comparison, a linear effects model was used (fitlm, Statistics Toolbox, Matlab, Mathworks Inc., Natick, MA); the dependent variable was defined as the tandem walk performance in VR+DOS, post bed rest, as well as post Shuttle and ISS missions. Statistical significance was determined using a 0.05 alpha level test.

## RESULTS

None of the subjects experienced stomach discomfort or nausea during the test sessions, although some subjects exhibited other signs of motion sickness, such as sweating, disorientation, and mild pallor. When performing EVA or IVA with 100% of their weight offloaded, many subjects reported slight vertigo and anxiousness about falling. When 62% of their body weight was offloaded, subjects reported increased difficulty performing basic functional tasks, such as walking or maneuvering to a standing position. Some subjects had feelings of disorientation and dizziness that persisted for up to 2 hours after they emerged from the VR+DOS condition.

The Percent of Correct Steps measured in each subject and condition is shown in Figure 5. Large variations in the Percent of Correct Steps was related to the subjects’ foot size, where they began the tandem walk on the foam surface, and the amount of stumbling during a trial. A repeated-measures ANOVA with 2 factors (visual conditions: VR, VR+DOS; body weight: 1 G, 0.38 G) indicated that the Percent of Correct Steps was significantly different in the VR and VR+DOS conditions [F(1,31) = 7.62, P = 0.01] and was significantly different in 1 G and 0.38 G [F(1,28) = 5.14, P = 0.03]. However, the interaction between the two factors was not statistically significant [F(1,28) = 0.61, P = 0.43]. Post hoc paired sample t-test indicated that the Percent of Correct Steps was significantly smaller in VR+DOS compared to VR for both gravity levels (offloaded body weights). This decrease was 46.6% in 1 G (P = 0.02) and 21.0% in 0.38 G (P = 0.02). In both visual conditions, the Percent of Correct Steps increased from 1 G to 0.38 G in 6 out of 8 subjects. This increase was significant in the VR+DOS condition (P = 0.04), but not in the VR condition (P = 0.18).

**Figure 5.**
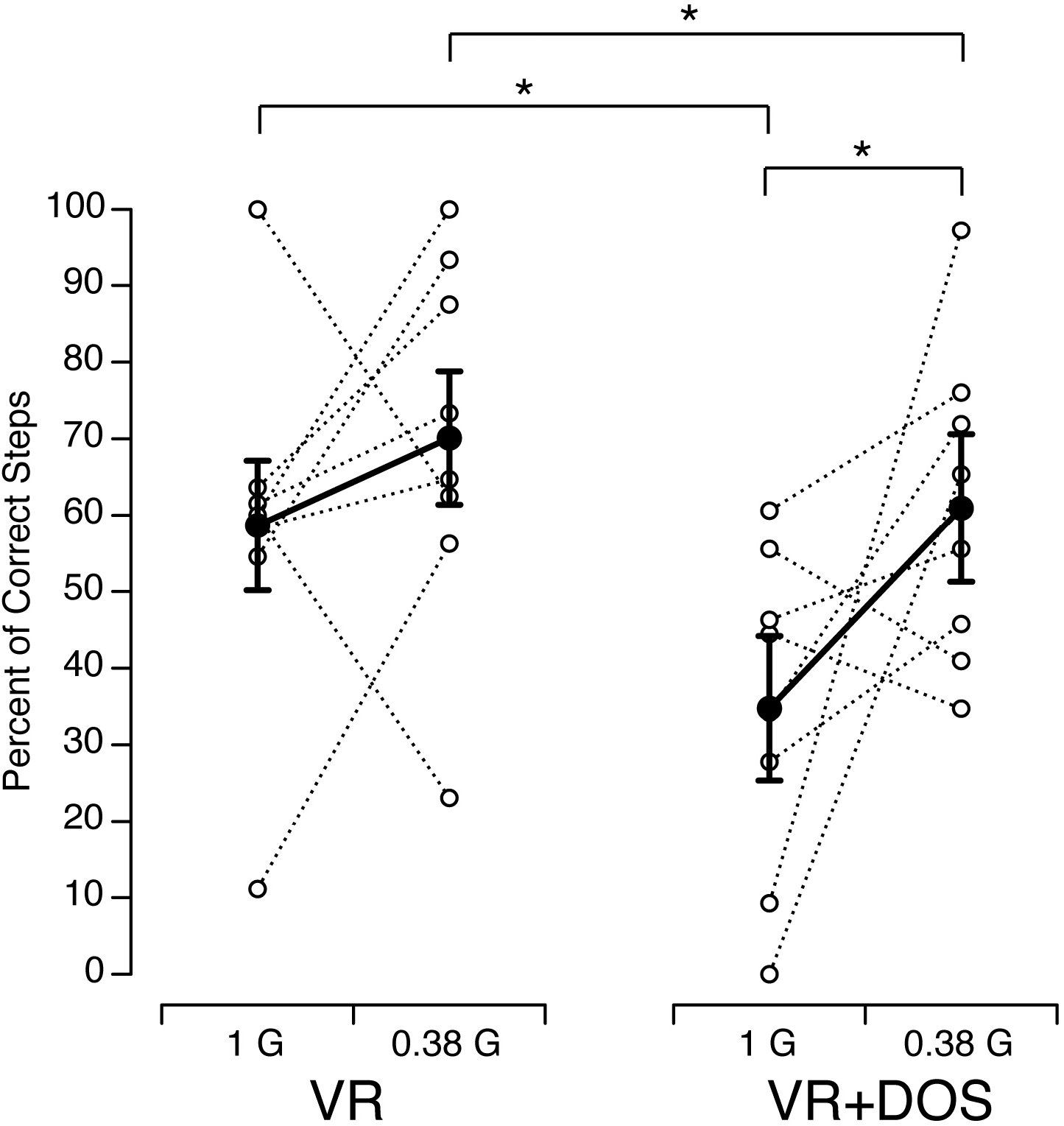
Percent of Correct Steps during the Tandem Walking Test in 8 subjects as a function of gravity level while viewing the virtual martian scene (VR) or the virtual martial scene with a superimposed optokinetic stimulation (VR+DOS). Open symbols show individual data; filled symbols show mean ± SD. * P < 0.05.

Figure 6 compares the Percent of Correct Steps in our study with those measured in astronauts after space missions and bed rest. The mean Percent of Correct Steps during tandem walking on a foam surface in VR+DOS in 1 G in our study (31.4 ± 17.1%; n=10) was not significantly different from the mean Percent of Correct Steps during tandem walking on a hard surface with the eyes closed immediately after a 2-week space flight (45.1 ± 20.4%; n = 7, P = 0.17) or a 70-day head down tilt bed rest (40.2 ± 21.4%; n = 10, P = 0.32), or one day after returning from a 6-month space flight (44.6 ± 19.7%; n = 13, P = 0.12).

**Figure 6.**
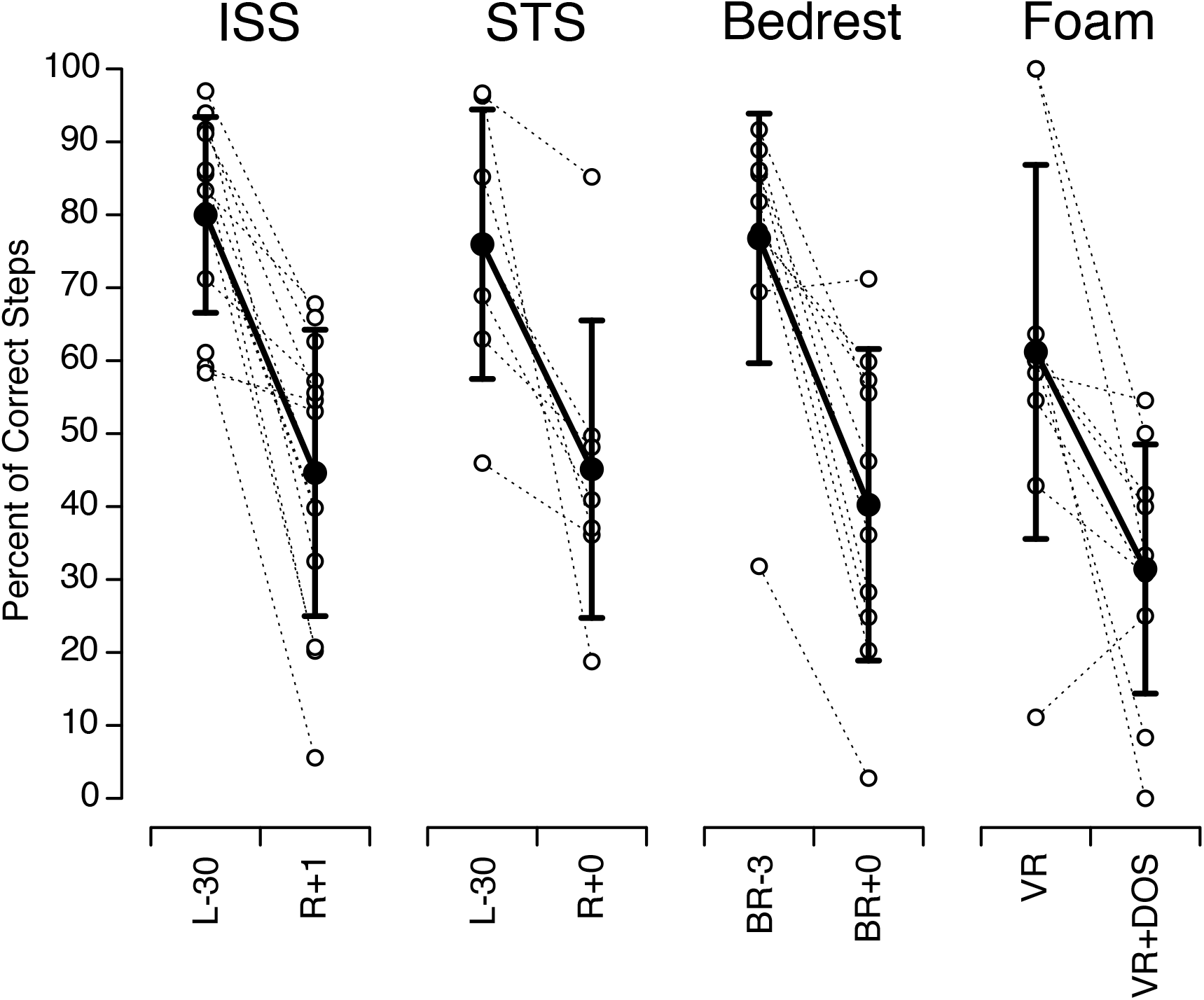
Comparison of the Percent of Correct Steps during tandem walk on a foam surface in our study (Foam) with the mean Percent of Correct Steps during tandem walk on a hard floor with the eyes closed in 13 astronauts before and after a 6-month stays on board the International Space Station (ISS) (adapted from Mulavara et al., 2018, Fig 5, p. 1970), in 7 astronauts before and after a 2-week spaceflight on board the Space Shuttle (STS) (adapted from Miller et al., 2018, Fig. 3, p. 811), and in 10 subjects before and after a 70-day bed rest in 6-deg head down tilt (adapted from Mulavara et al., 2018, Fig 5, p. 1970). Open symbols show individual data; filled symbols show mean ± SD. L-: day before flight; R+: day after flight; BR-: day before bed rest; BR+: day after bed rest.

## DISCUSSION

In this study, we investigated how simulated martian gravity and visual disorientation affect tandem walk performance. Results indicate that tandem walk performance was altered when subjects were viewing a visual disorientation superimposed to a virtual reality scene (VR+DOS). However in this visual condition, the tandem walk performance was better in 0.38 G than in 1 G. Results also show that the performance of our subjects during the Tandem Walking Test on a foam flooring in VR+DOS was comparable the performance of astronauts during tandem walk on a hard flooring with the eyes closed after spaceflight (Miller et al. 2018, Mulavara et al. 2018).

The purpose of the Tandem Walking Test was to assess changes in dynamic balance control. Dynamic balance during walking is controlled by motor neurons that regulate muscle activity, and by sensory inputs from vestibular, visual, and somatosensory systems, as well as a central prediction of the afferent signals (Woollacott & Shumway-Cook, 2002).

### Role of the Vestibular System

The Tandem Walking Test is a reliable indicator of vestibular driven ataxia (Fregly et al., 1972). Patients with vestibular impairments have impaired tandem walking performance compared to healthy subjects (Cohen et al., 2012). In addition, it has been shown that tandem walking performance declines with age (Speers et al., 1998; Vereeck et al., 2008).

When walking, head stabilization contributes to dynamic balance because head movements in pitch compensate for the vertical trunk translation that occurs during each step of the gait cycle (Pozzo et al., 1990). In addition to head stabilization, the vestibulo-ocular reflex compensates for the head movements to stabilize gaze in space. This coordinated strategy between the motion of the eye, head, and trunk plays a central role in maintaining clear vision during natural body movements (Bloomberg et al., 1997). The otolith organs of the vestibular system participate in the gain of the vestibulo-ocular reflex during movement of the head in pitch and in yaw. Individuals with vestibular deficits also restrict their head movements during locomotion (Grossman & Leigh, 1990).

Astronauts returning from spaceflight and walking through an obstacle course on the same compliant foam surface as in our ground-based study exhibited altered locomotor function, with a median 48% increase in the time to complete the course (Mulavara et al., 2010). Decrements in computerized dynamic posturography were also seen in the same crewmembers (Wood et al., 2015). After 2-week spaceflight, the ability to stand on rails with the feet in tandem (Tandem Romberg Test) was reduced to 40%-90% in Space Shuttle crewmembers (Homick & Reschke 1977). However, a decrease in tandem walk performance, similar to that observed in returning astronauts, was also observed after 60-d and 70-d head-down tilt bed rest in ground-based subjects (Macaulay et al., 2016; Miller et al., 2018; Mulavara et al., 2018). During bed rest and in our study, the vestibular system was still exposed to the 1 G gravitational acceleration, therefore the decrement in tandem walk performance in all of these studies cannot be solely attributed to decreased vestibular function (Mulavara et al., 2008).

### Role of the Visual and Somatosensory Systems

Viewing the VR+DOS scene altered the contribution of the vestibulo-ocular reflex for stabilizing gaze. This result is in agreement with computerized dynamic posturography studies showing greater imbalance during optokinetic stimulation (Van Ombergen et al., 2016) and when the visual input is stabilized relative to the head (Nashner et al. 1982) than in darkness. Ground-based studies have also shown that older adults use vision to a greater degree to control head stabilization than do young adults (Keshner et al. 1995; Cromwell et al., 2002). The increased reliance on vision by older adults has been attributed to a decrease in vestibular and proprioceptive function with age (Peterka & Black, 1990).

The increased tandem walk performance in ARGOS in 0.38 G could be due to an increase in the role of somatosensory inputs in controlling dynamic balance when the body is unloaded. ARGOS reduces body weight bearing, which modifies the inputs from the receptors in the skin, muscles, bones, joints, and internal organs. Body unloading reduces the inputs from the mechanoreceptors in the sole of the feet. However, other skin areas in contact with the harness provide inputs for the control of dynamic balance. Muscular tone in the leg postural muscles is also reduced due to body unloading, and the efferent signals generated by the motor system are different.

The improved performance of tandem walk when the body weight was unloaded in ARGOS compared to 1 G could be because the body harness in ARGOS helps minimize body sway, thus improving postural stability and gait. The reduced ground reaction force when body weight is unloaded also contributes to an increased cone of stability. The limits of stability are defined by the angles in which the body’s center of gravity moves past the base of support, and when the limits are reached a correction is required (e.g., a step) to maintain balance. Less ground force reaction allows further deviation from the base of support before a correction is required, thus less ground force facilitates postural balance and gait. Ground-based studies using isokinetic testing system have shown that proprioception was significantly correlated with the limits of stability (Wang et al., 2016).

### Spaceflight Analog

Recent investigations by our group showed a significant reduction in the Percent of Correct Steps during the Tandem Walking Test with the eyes closed performed by astronauts immediately after short- or long-duration spaceflight (Miller at al., 2018; Mulavara et al., 2018). After astronauts return from a spaceflight, the width of their leg placement is exaggerated, they take small steps of irregular length, and they fail to maintain their intended path (Bloomberg et al., 1997). Astronauts often experience oscillopsia when they move their head after returning from space, which contributes to disrupted locomotion (Reschke et al., 2017). The results of the present study indicate that an exaggerated motion of the visual scene during head movements (VR+DOS) in 1 G impairs walking performance on a compliant foam floor in the same manner as in astronauts with the eyes closed immediately after spaceflight. The combination of a medium-density foam surface, VR, and optokinetic stimulation could therefore be used as an analog to evaluate the effectiveness of countermeasures for mitigating walking performance after spaceflight. The system could be improved, however, by adding galvanic vestibular stimulation triggered with head movements to alter the contribution of the vestibular inputs, as is the case after spaceflight (Wood et al., 2011). Additionally, further work is required to determine if this analog environment could be used to expose crewmembers to the type of visual-vestibular deficits they will experience after landing, and determine if preflight training can improve postflight walking performance.

## ACKNOWLEDGMENTS

The author thank Kerry George for her editorial recommendations. The authors would like to thank Anjul Patney from NVIDIA for aide with the OpenVR integration and Falcor source code. The authors would also like to thank Adriel Arias for help with modeling the Mars terrain. This work was completed with the NVIDA Falcor engine (Benty et al., 2017). This work was supported by the National Aeronautics and Space Administration (NASA).

